# Mantis: flexible and consensus-driven genome annotation

**DOI:** 10.1101/2020.11.02.360933

**Authors:** Pedro Queirós, Francesco Delogu, Oskar Hickl, Patrick May, Paul Wilmes

## Abstract

**Background:** The past decades have seen a rapid development of the (meta-)omics fields, producing an unprecedented amount of data. Through the use of well-characterized datasets we can infer the role of previously functionally unannotated proteins from single organisms and consortia. In this context, protein function annotation allows the identification of regions of interest (i.e. domains) in protein sequences and the assignment of biological functions. Despite the existence of numerous tools, some challenges remain, specifically in terms of speed, flexibility, and reproducibility. In the era of big data it also becomes increasingly important to cease limiting our findings to a single reference, coalescing knowledge from different data sources, thus overcoming some limitations in overly relying on computationally generated data.

**Results:** We implemented a protein annotation tool - Mantis, which uses text mining to integrate knowledge from multiple reference data sources into a single consensus-driven output. Mantis is flexible, allowing for total customization of the reference data used, adaptable, and reproducible across different research goals and user environments. We implemented a depth-first search algorithm for domain-specific annotation, which led to an average 0.038 increase in precision when compared to sequence-wide annotation. Mantis is fast, annotating an average genome in 25-40 minutes, whilst also outputting high-quality annotations (average coverage 81.4%, average precision 0.892).

**Conclusions:** Mantis is a protein function annotation tool that produces high-quality consensus-driven protein annotations. It is easy to set up, customize, and use, scaling from single genomes to large metagenomes. Mantis is available under the MIT license available at https://github.com/PedroMTQ/mantis.

## Background

Life, on a cellular scale, is, in essence, the activity and the interaction of a plethora of different molecules, among which proteins cover the primary role of carrying out the vast majority of processes. A major task in understanding how biology works is to be able to properly recognise its actors (e.g. the proteins) and put them into context. The past decades have seen the development of the (meta-)omics fields, unlocking an unprecedented amount of data and deepening our understanding in several fields of biology [66]. Alongside the evolution of the technologies and the increase in data volume, the identification of proteins transitioned from purely experimental techniques (e.g. chemical essays, spectroscopy, etc.) toward the computational-based sequence analysis thanks to the discovery of the relationship between conservation of proteins’ functions and sequences [82]. Therefore, the current challenges are to make use of the vast number of protein sequences and annotations available and to link new protein sequences to the previously established knowledge. High-throughput methods, such as next-generation sequencing, are able to produce a large amount of data which then needs to be analysed and interpreted. One of the ways to make sense of this data is through protein function annotation (PFA), which is the identification of regions of interest (domains) in a sequence and assignment of biological function(s) to these regions. This strategy has proven effective in both, the study of single organisms as well as consortia [5, 13, 31, 26, 40, 47]. Function prediction is based on reference data, i.e. transferring the function from protein X to the unknown protein Y if they are highly similar [82]. Different approaches may be used, the most common being to compare an unknown protein sequence to a reference database composed of well-studied and annotated proteins (homology-based methods) [71, 79, 10, 70, 65, 30]; other methods may infer function through the use of machine learning [71, 63], protein networks [88, 72], protein structure [15], or genomic context-based techniques [44], but these will not be covered in this paper. For sequence alignment BLAST [2] or Diamond [11] are commonly used, whereas, for HMM profiles, HMMER [62] is most widely used. In PFA, these tools are often integrated into larger pipelines to provide enhanced output interpretability, workflow automation, and parallelization. [65, 30, 7, 32]. Some PFA tools target specific taxa [39], others are designed with large-scale omics analysis in mind [85, 42, 34]; indeed, each PFA tool is designed to cater to its niche research topic. While experimental validation remains the gold standard, PFA, despite its many shortcomings [51], is an increasingly valuable strategy that aims to tackle the progressively more difficult task of making sense of the large quantities of data continuously generated.

We reviewed the implementation of three widely used PFA tools [65, 32, 30] and observed that the processing of candidate annotations (i.e. sequences or HMM profiles which are highly similar to the query sequence) is done by capturing only the most significant candidate between the references (“best prediction only”, herein after called **BPO**); to our knowledge, this is the most common method. This classic PFA approach works well for single-domain proteins, but multidomain proteins may have multiple putative predictions [86, 17, 37], whose location in the sequence may or may not overlap. This selection criterion may potentially lead to missing annotations and is therefore not suitable in complex PFA scenarios. To tackle this problem, domain-specific PFA is necessary. A simple approach, previously discussed in Yeats et al.[86], would be to order the predictions by their significance and iteratively add the most significant one, as long as it does not overlap with the already added predictions (henceforth referred to as the **heuristic** algorithm). Due to the biased selection of the first prediction, this algorithm does not guarantee an optimal solution (e.g. a protein sequence may have multiple similarly significant predictions). It has been previously shown that incorporating prediction significance and length may produce better results [76]. We implemented a Depth-First Search (**DFS**) algorithm that improves on the previous approaches.

The selection of reference HMMs is also extremely important, as PFA is based on the available reference data. Whilst sing unspecific HMMs to annotate a taxonomically classified sample may result in a fair amount of true-positives (correct annotations), depending on how strict the confidence threshold is, it may also increase the false-positives (over-annotation, due to low confidence threshold) or false-negatives (under-annotation, due to high confidence threshold) [64]. Using taxa-specific HMMs (TSHMM) rather than unspecific HMMs should, in principle, provide better annotations on a taxonomically classified sample, a feature that is already integrated into some PFA tools such as eggNOG-mapper [30] and RAST [7]. In essence, TSHMMs-based annotation limits the available search space, which may have both positive and negative consequences. Since the search space is more specific, the annotations produced should be of higher quality; however, this high specificity of the TSHMM could also lead to under-annotation (incomplete reference TSHMMs) or mis-annotations (low-quality reference TSHMM) [19]. This underlines the necessity to use specific (e.g. TSHMMs) and unspecific HMMs in a complementary manner. In this regard, the use of multiple reference datasets remains a challenging aspect of PFA, and, with multiple high-quality reference datasets available, it is increasingly important to be able to coalesce knowledge from different sources. Whilst some PFA tools do allow for the use of multiple reference data sources, either as a separate [32] or a unified [30, 4] database, it is still challenging to dynamically integrate multiple data sources.

When using multiple high-quality reference datasets, the most common, and simplest, approach is to consider the output from each reference data source independently (e.g. [32]). However, by doing so we overlook that many sources can overlap and/or complement each other. Commonly this is compensated via manual curation, which is feasible only for a limited number of annotations. An automated approach would be to assume only the most significant annotation source for any given sequence, disregarding other sources, which may result in vast losses of potentially valid and complementary information. As this is not desirable, the challenge is both in deciding which source(s) provide the best annotation as well as identifying complementary annotations. A straightforward approach to solve this issue would be to assume that the sequence annotations are complementary when they share a database identifier, for example:

1. Function: “Responsible for glucose degradation”; Identifiers: K00844,EC:2.7.1.1,**PF03727**
2. Function: “Responsible for glucose degradation”; Identifiers: P52789,**PF03727**,IPR022673
3. Function: “Protein is an enzyme and it is responsible for the degradation of glucose”; Identifiers: HXK2_HUMAN

We can observe that the annotations (i) and (ii) share the database identifier **PF03727**, and thus it can be concluded that these are complementary annotations. If we were only to select the first annotation, we would ignore potentially useful information (identifiers P52789 and IPR022673). However, it may be the case that no identifiers are shared between the different annotations, for example:

1. Function: **”Responsible for glucose degradation”**; Identifiers: K00844,EC:2.7.1.1
2. Function: **”Responsible for glucose degradation”**; Identifiers: P52789,IPR022673
3. Function: “Protein is an enzyme and it is responsible for the degradation of glucose”; Identifiers: HXK2_HUMAN

We can observe that even though the annotations (i) and (ii no longer share an identifier, they still have the same function “**Responsible for glucose degradation** “. Humans can quickly surmise that these annotations are the same as they share the same function description. Should the descriptions be identical or very similar, a machine could achieve the same conclusion with relative ease. However, in our experience, these free-text descriptions are often moderately or heavily dissimilar [35, 69], with only a few keywords allowing us to ascertain they are indeed the same. This then makes it more difficult to use multiple reference sources. For example:

1. Function: “Responsible for **glucose degradation** “; Identifiers: K00844,EC:2.7.1.1
2. Function: “Protein is an enzyme and it is responsible for the **breakdown** of **glucose**”; Identifiers: HXK2_HUMAN

In such a scenario, someone trained in a biology-related field can quickly identify the most important words (”degradation”/”breakdown” and “glucose”) in both sentences and conclude both annotations point to the same biological function. The challenge is now to enable a machine, deprived of any intellect and intuition, to eliminate confounders (very common words, e.g. “the”), identify keywords and their potential synonyms, and reach the same conclusion. A possible strategy is to use text mining, this is the process of exploring and analysing large amounts of unstructured text data aided by software, identifying potential concepts, patterns, topics, keywords, and other attributes in the data [20]. Text mining has been previously used with biological data [80, 49, 87, 68, 27], and even more specifically with regards to gene ontologies [9, 48, 38, 14, 16, 36] and PFA [87]. However, to our knowledge, there is no tool for the dynamic generation of a consensus from multiple protein annotations. This paper solves the problem of scaling the integration of different annotation sources, integrating a compact and flexible text mining strategy. We implemented a two-fold approach to build a consensus annotation, first by checking for any intersecting annotation identifiers and second by evaluating how similar the free-text annotation descriptions are. This approach addresses three very relevant issues with PFA [64, 19, 52, 18]: over-annotation (through the use of overlapping but independent sources, thus obtaining a more reliable final annotation); under-annotation (through the use of multiple reference sources, which implicitly leads to a wider search space); and elimination of redundancy (through the creation of a consensus-driven annotation).

Another challenge in PFA is the lack of flexibility of some tools, as these are often intrinsically connected to their in-house generated reference data, and therefore very hard to customize. In contrast, we developed a tool that while offering high-quality unspecific and specific HMMs, it is independent of its reference data, thus being customizable and allowing dynamic integration of new data sources.

We hereby present Mantis, a Python-based PFA tool that overcomes the previously presented issues, producing high-quality annotations with the integration of multiple domains and multiple reference datasets. Mantis automatically downloads and compiles several high-quality reference sources, and is able to make the most efficient use of the available hardware through parallelized execution. Mantis is independent of any of the default reference datasets, being flexible and customizable, resulting in a versatile and reproducible tool that overcomes the challenge of high-throughput protein annotation coming from the many genome and metagenome sequencing projects.

## Mantis

Mantis is available at https://github.com/PedroMTQ/mantis [60] and its workflow (see **Figure 1**) consists of 6 main steps: i) sample pre-processing; ii) HMM profile-based homology search against each reference dataset; iii) intra-HMM reference hits processing; iv) metadata integration; v) inter-HMM reference hits processing; and vi) consensus generation. During sample pre-processing, the sample(s) is/are split into chunks so that the homology search can be parallelized. During homology search, query sequences are searched against all available reference datasets using HMMER. During intra-HMM hits processing the DFS algorithm is used to generate and select the best combination of hits per HMM source. During metadata integration, metadata (annotation description and identifiers) is added to the respective hits. During inter-HMM hits processing the DFS algorithm is again used to generate all the combinations of hits for all HMM sources. During consensus generation all the combinations of hits are expanded and intersected (if possible), the best consensus combination of hits is then selected for each query sequence (see **Methods** for a detailed description of all these steps). Alongside Mantis, We developed a standalone tool [53] that allows the use of multiple reference datasets through the generation of a consensus annotation. As we have shown in the supplement “Defining a similarity threshold”, this tool has high specificity, thus, in the context of Mantis, it allows for the correct identification of similar free-text annotation descriptions.

**Figure 1:**
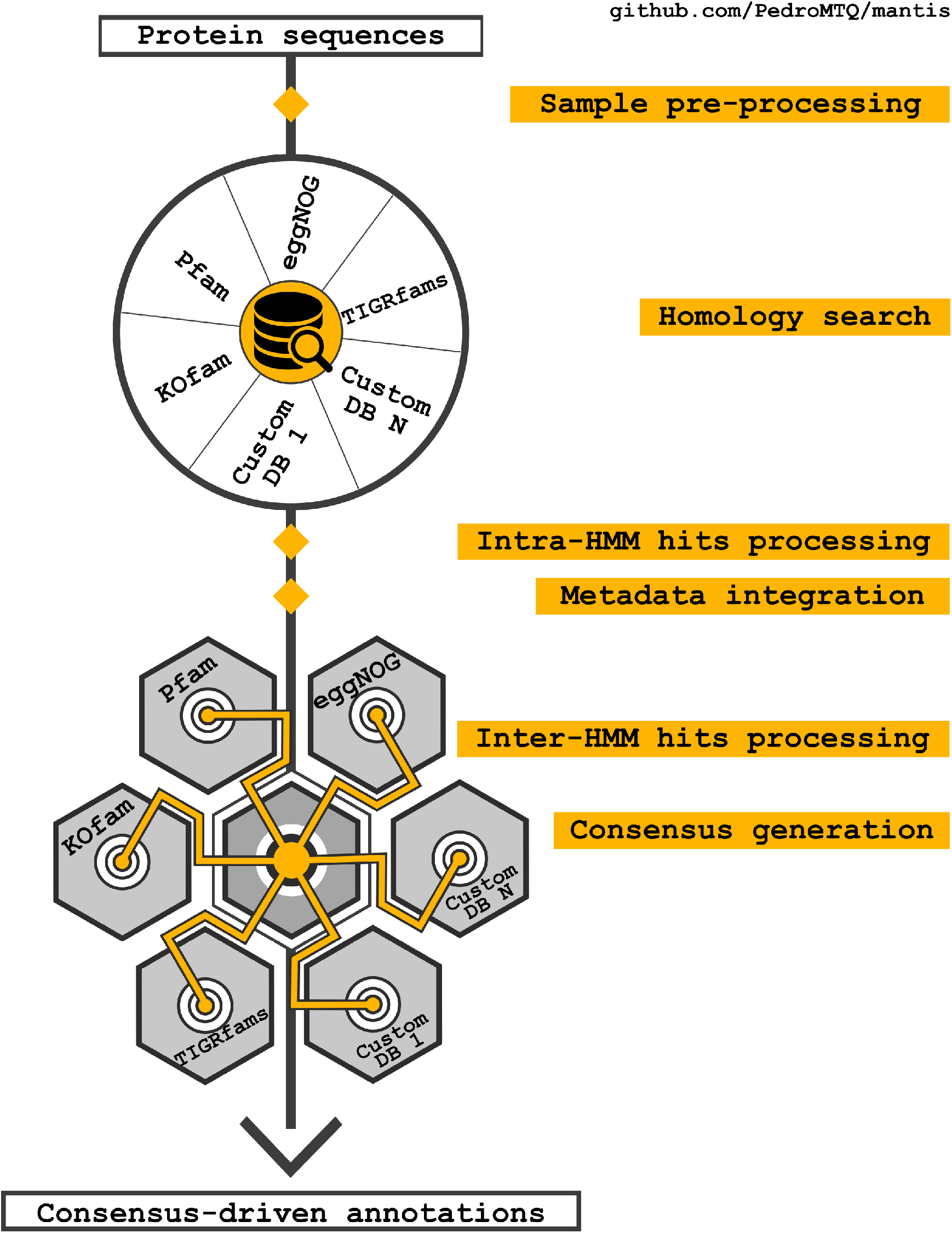
Overview of the Mantis workflow. KOfam [3], Pfam [21], eggNOG [29], and TIGRfams [22] are the reference HMMs currently used in Mantis. CustomDB can be any HMM library provided by the user.

## Analysis

To analyse and validate the performance of Mantis, we performed several in-silico experiments: As an initial quality control analysis, we started by annotating a reference dataset from Swiss-Prot such that we are aware of the optimal execution parameters. In this context, we evaluated the impact of the e-value threshold, the hit processing algorithm, and the contribution of each reference dataset to the final output. After this initial quality control analysis, we annotated several sequenced organisms, with and without using TSHMMs, thus evaluating the impact of using taxa-resolved reference datasets. Finally, we compared Mantis against eggNOG-mapper [30], a similar PFA tool known for producing taxa-resolved annotations. The sample description is available in “ **Sample selection**”.

### Initial quality control

#### Function assignment e-value threshold

It is known that the e-value threshold directly affects annotation quality, however, no gold-standard threshold exists [76]. Depending on the size, quality, and specificity of the reference data source, we may use more or less stringent thresholds. It is therefore essential to test annotation quality with different thresholds. As such we tested different static e-value thresholds as well a dynamic threshold, which has been described in “**Establishing a test environment**. Testing different e-value thresholds”. As seen in supplemental **Table 1**, being more permissive (by using a higher e-value threshold e.g. 1*e*^−3^) resulted in a higher annotation coverage and a higher precision. This is due to Mantis’ internal quality control in the form of the DFS hit-processing algorithm. Being too strict with the e-value threshold constricts the available solution space, resulting in a lower precision. Based on these results the default e-value threshold was set to 1*e*^−3^.

#### Impact of sample selection and hit processing methods on precision

Testing exclusively against well-annotated organisms is a recurring issue with protein annotation benchmarking, resulting in the re-annotation of sequences already present in the reference datasets used, leading to a biased annotation quality evaluation. To avoid this bias, we downloaded all Swiss-Prot protein entries (as of 2020/04/14) and selected entries by their creation date such that we have four samples that contain protein entries created in different date ranges (2010-2020, 2015- 2020, 2018-2020, and 2020). Since the default reference datasets were compiled before 2020, we ensure that the sample with protein entries from 2020 is not in the reference datasets used, thus guaranteeing that potential annotations are due to true sequence homology. We annotated these samples using three different hit processing algorithms (DFS, heuristic, and BPO), determining the impact of each on precision.

As seen in **Figure 2**, precision decreased as the sample was restricted to data from more recent years. When comparing the hit processing algorithms we found that the DFS algorithm consistently outperformed the other algorithms, with an average precision 0.038 and 0.013 higher than the BPO and heuristic algorithms respectively. In addition, the precision difference between the multiple predictions algorithms (DFS and heuristic) and the single predictions algorithm (BPO) increased as the entries in a sample were restricted to more recent years. Further details can be found in the supplemental **Table 2**.

**Figure 2:**
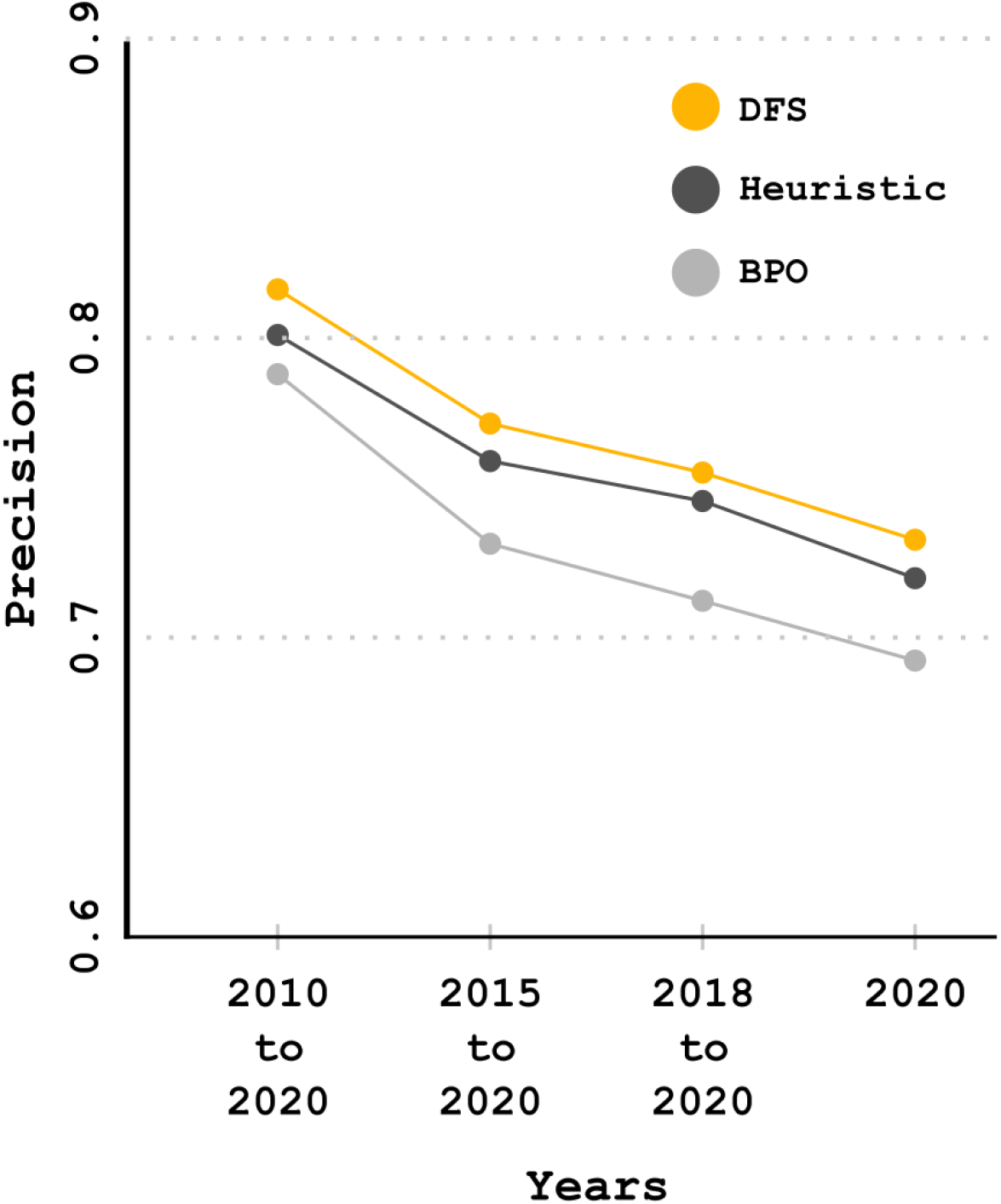
Annotation precision per hit-processing algorithm and Swiss-Prot sample. The DFS algorithm outperforms the other algorithms.

#### Contribution of the different reference datasets

We also analysed the contribution of each reference dataset to the output annotation for the biggest Swiss-Prot sample (2010-2020). Pfam was present in 17.6% of the annotations, KOfam in 32.5%, eggNOG in 37.9%, and TIGRfam in 10.1%.

#### Quality control against sequenced organisms

As a secondary quality control, we annotated several sequenced organisms (for more details see the supplemental **Table 3**) with and without TSHMMs, to assess the impact on precision when using taxa-resolved references. We also evaluated the impact of the different hit processing algorithms on this dataset.

As seen in **Figure 3**, well-studied organisms, such as *Escherichia coli* and *Saccharomyces cerevisiae*, had better annotations, especially when applying TSHMMs, unlike poorly described organisms. The average precision gain with TSHMMs was 0.029. With TSHMMs, the DFS algorithm had, on average, 0.0001 and 0.0116 higher precision than the BPO and heuristic algorithms respectively. Without TSHMMs, the DFS algorithm had, on average, 0.0126 and 0.0189 higher precision than the BPO and heuristic algorithms respectively. The respective results by organism and algorithm can be seen in detail in the supplemental **Table 4**.

**Figure 3:**
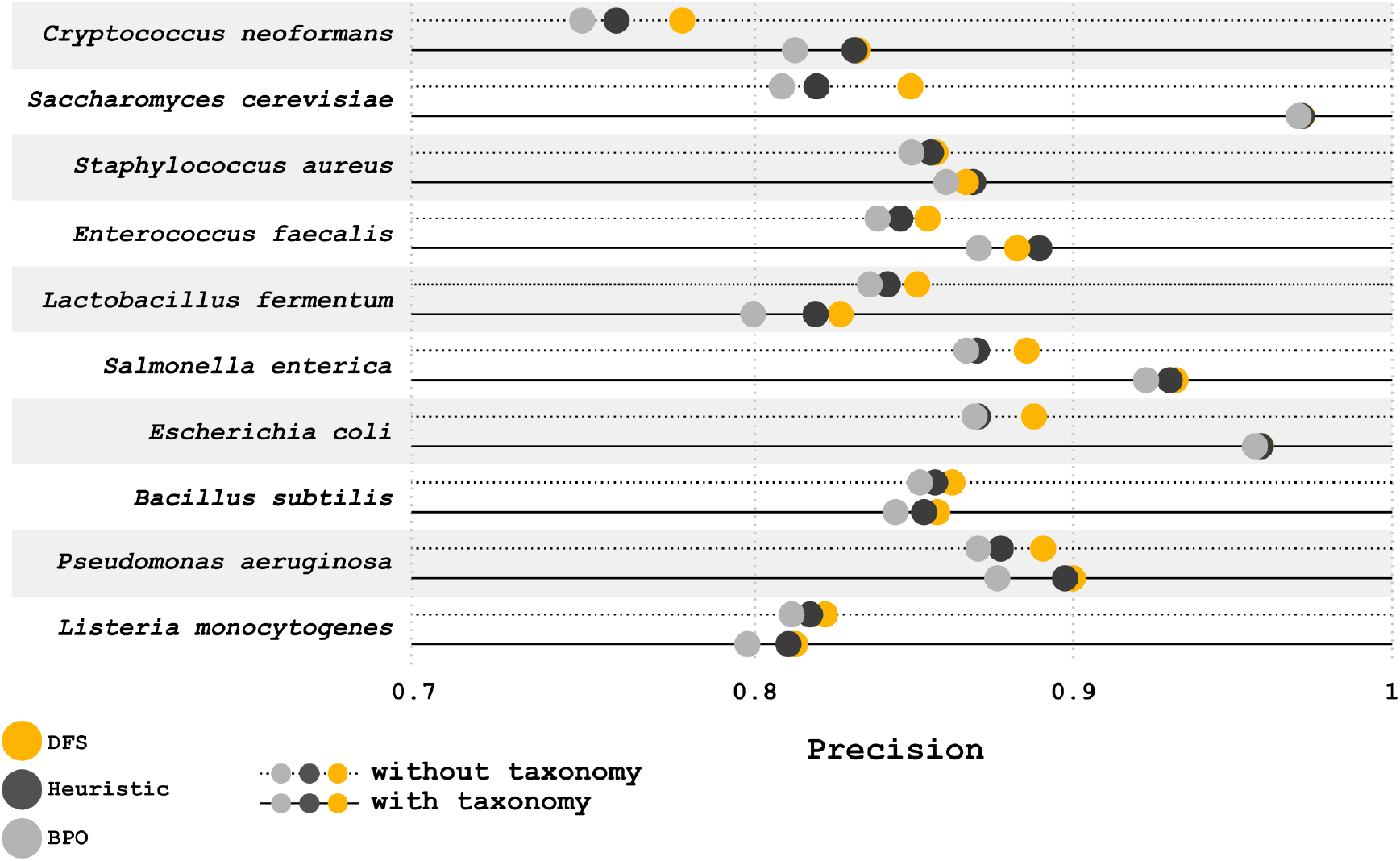
Annotation precision per hit-processing algorithm and organism, with and without using taxonomy information. Precision was higher for well-studied organisms, TSHMMs also performed better with these organisms.

#### Comparison between Mantis and eggNOG-mapper

The sequenced organisms enumerated in the supplemental **Table 3** were annotated with Mantis and eggNOG-mapper. To evaluate the added value of using multiple reference datasets against exclusively using eggNOG we also assessed Mantis’ precision using different reference datasets: (i) Mantis with default data sources and with taxonomy information; (ii) Mantis with default data sources except for eggNOG’s data (iii) Mantis exclusively with eggNOG’s data and with taxonomy information; and (iv) Mantis exclusively with eggNOG’s data and without taxonomy information.

On average, Mantis’ default setting (including eggNOG as a reference dataset) and eggNOGmapper had a precision of 0.89 and 0.78, respectively, and, as seen in **Figure 4**, Mantis’ default setting outperformed eggNOG-mapper for all the benchmarked organisms. Comparatively, Mantis’ default setting had, on average, 8.26 % more true-positives (TP) than eggNOG-mapper, 8.77% fewer false-positives (FP), 0.76% less coverage, and 0.24% fewer potentially new annotations. To surmise, despite minimal losses in coverage, the default Mantis execution had a precision 0.11 higher than eggNOG-mapper.

**Figure 4:**
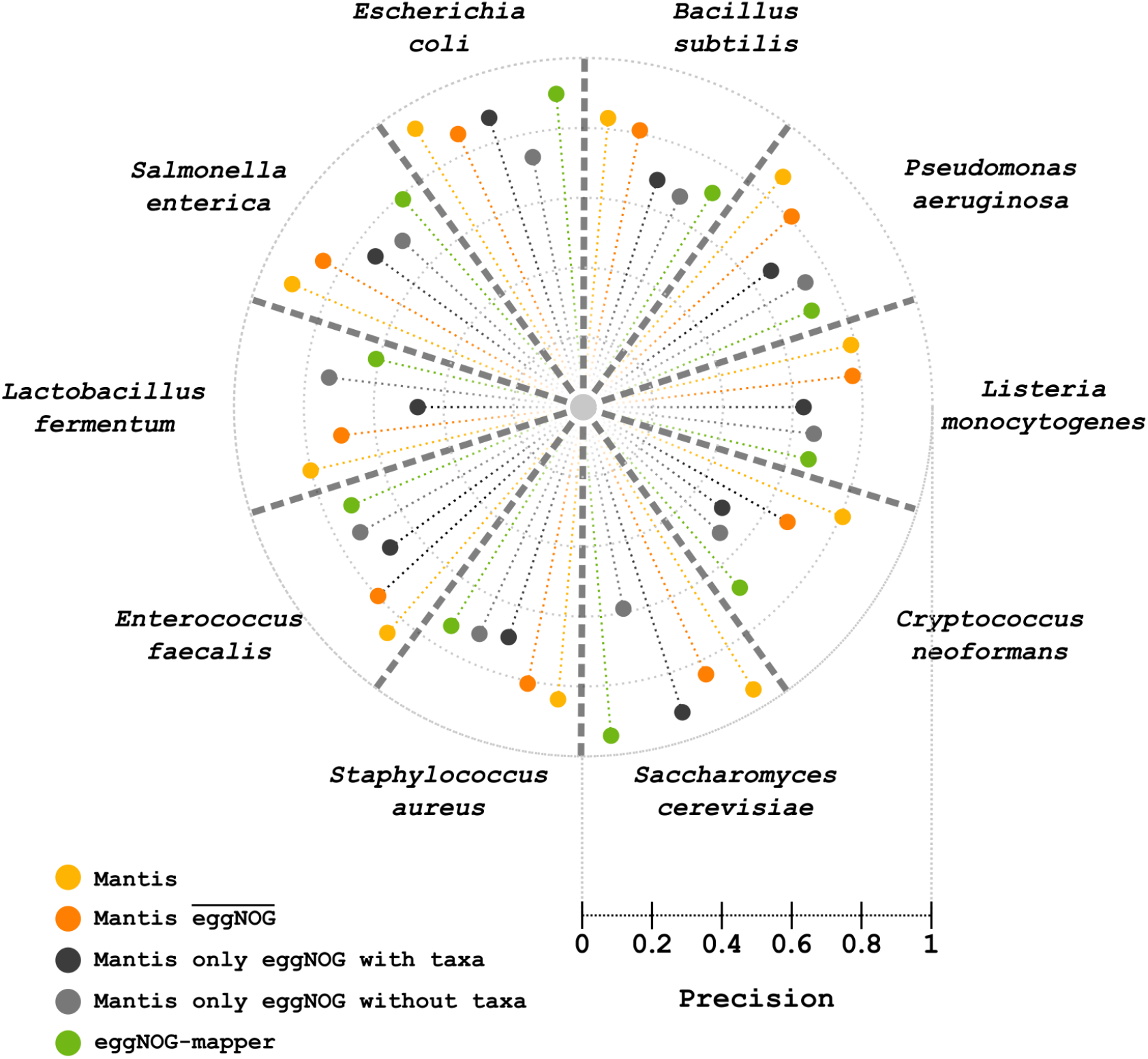
Annotation precision of Mantis and eggNOG-mapper using different reference datasets. Each slice represents to an organism and contains the attained annotation precision between the different conditions

Mantis’ setting without eggNOG had an average precision 0.06 higher than eggNOG-mapper and an average precision 0.05 lower than Mantis’ default setting. When only eggNOG’s data (with and without TSHMMs) is used by Mantis, eggNOG-mapper outperforms it. When TSHMMs were not used, eggNOG-mapper had a 0.055 higher precision than Mantis. When eggNOG’s TSHMMs were used, eggNOG-mapper had a precision 0.045 higher than Mantis. Further details are available in the supplemental **Table 5**. On average, Mantis’ default setting took 37 minutes, Mantis without eggNOG 18 minutes, and eggNOG-mapper 3 hours and 22 minutes. Average seconds taken to annotate a sequence were also compared: on average we found that Mantis was 6.6 times faster than eggNOG-mapper; Mantis without eggNOG ran 14.7 times faster than eggNOG-mapper and 2.2 times faster than the default Mantis execution. Further runtime benchmarking can be found in supplements **”Benchmarking annotation efficiency”**.

## Discussion

We herein presented Mantis, a PFA tool that produces high-quality annotation and is easily installed and integrated into other bioinformatic workflows. As shown in the **Accessibility and Scaling**, a conda environment and an automated reference datasets download are provided. In addition, Mantis accepts several formats as input (i.e. protein fasta file, tsv file with paths, directories, or compressed archives), outputting easy to parse tsv files. Mantis also addresses some major challenges in PFA, such as flexibility, speed, the integration of multiple reference datasets, and use of domain-specific annotations. Mantis uses a well-established homology-based method and produces high-quality consensus-driven annotations by relying on the synergy between multiple reference datasets and improved hit processing algorithms. Mantis’ default execution resulted in an average coverage of 81.3% and average precision of 0.89 when annotating a widely used, well characterized microbial community standard, as seen in **Figure 4**.

Well-curated and commonly used resources were chosen as the default reference datasets for Mantis, containing both unspecific and specific reference datasets (e.g. taxa-specific). As we have shown no single reference dataset accounted for the majority of the annotations, each offering both unique and overlapping insight into protein function, thus confirming their synergy and redundancy.

Multiple reference datasets are integrated through a consensus-driven approach, which Mantis uses as an additional quality control step, as well as a means to automatically incorporate a broader variety of identifiers. Our text mining technique eliminates confounders and identifies the most important keywords in the annotation descriptions, thus making similarity comparisons between descriptions possible. Our method is flexible and adaptable towards other lexicons and pipelines and is available as a standalone tool [53]. Our method would benefit from a biological synonym lexicon, indeed, while non-biological words are easy to deal with (e.g. using Wordnet’s lexicon [84]), the same cannot be said for biological terms. Biological lexicons exist [75], but are often behind pay-walls, which reduces the potential use for open source projects trying to follow FAIR principles [83]. This limitation may be worked on in further versions of Mantis by applying vocabulary expansion methods such as the method described by Slater et al. [68]. Our text mining technique may also benefit from the implementation of a tagger specific to biomedical text, such as Hunflair [81], which we will test and implement in a future version of Mantis.

We have additionally implemented two algorithms for domain-specific homologs search (DFS and heuristic as back-up). **Figure 5** shows how dependent homolog selection is on the hit-processing algorithm used. We have not only shown that these algorithms are more precise when annotating previously described protein sequences but that their impact on precision significantly increased when annotating uncharacterized protein sequences (e.g. average precision gain with DFS and BPO algorithms in the Swiss-Prot samples 2010-2020 and 2020 were 0.028 and 0.040, respectively). We hypothesize that for the latter, homology search is not capable of finding whole-sequence homologs, finding however multiple domains that partially constitute the protein sequence. As such, we hypothesize that by increasing the resolution (sequence homology to domain homology) of homology-based reference datasets domain-specific algorithms may become increasingly useful. We think this would be especially important when annotating protein sequences without well-described homologs, but that contain previously characterized conserved protein domains.

**Figure 5:**
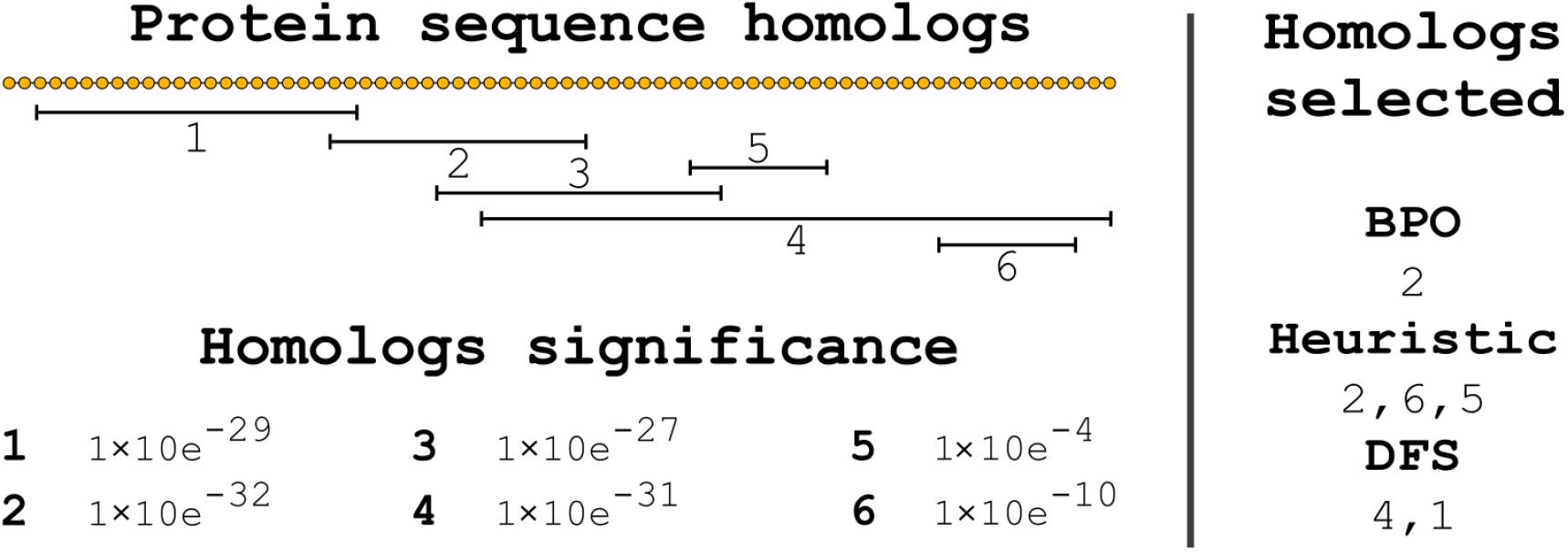
Homologs selection per hit processing algorithm. PFA is entirely dependent on the algorithm used to select the hits. For instance, BPO only selects the most significant hit (2) even though hit 4 is wider and has almost the same significance. The heuristic method instead initially selects the most significant hit (2) which then restricts the hits available for selection (5, 6). The DFS algorithm generates all possible combinations of hits and then selects the one that is both very significant and with large sequence coverage.

In **Figure 6.A** we can observe that the current query sequence is already used to generate the HMM profiles) in the reference datasets, thus matching with the HMM profile that contains it, and finally being annotated as a “coagulation factor”. Such a scenario is common when annotating well-described organisms (e.g. *Escherichia coli*). It may however be that the query sequence (in the scenario **Figure 6.B** is not in the reference dataset (which is common in non-model organisms or niche metagenomic samples), thus partially matching with several HMMs, which may correspond to multiple domains (depending on the resolution of the reference datasets). Unlike the BPO algorithm, the heuristic and DFS algorithms are able to incorporate multiple homologs, and while these may not be enough to determine the biological function of a protein, they still provide a better biological context than a single annotation. Further improvements in annotation quality may also require the use of motif-based and/or genomic context-based (e.g. operon context information, co-expression, and subsystems) methods such as those described by Sigrist et al.[67], Mooney et al. [43], Mavromatis et al.[41], Overbeek et al.[44], and Hannigan et al.[24]. Nevertheless, the significantly higher precision seen when comparing the DFS and BPO algorithms (as seen in supplemental **Table 2**) highlights the need to adopt better hit processing methods, especially for non-model organisms. With samples ranging from thousands to hundreds of thousands (or more) of protein sequences, sub-optimal hit processing algorithms may cascade into unnoticeable pitfalls in downstream data analysis (e.g. accumulation of incomplete or low quality genome annotation which may in turn lead to false biological interpretations).

**Figure 6:**
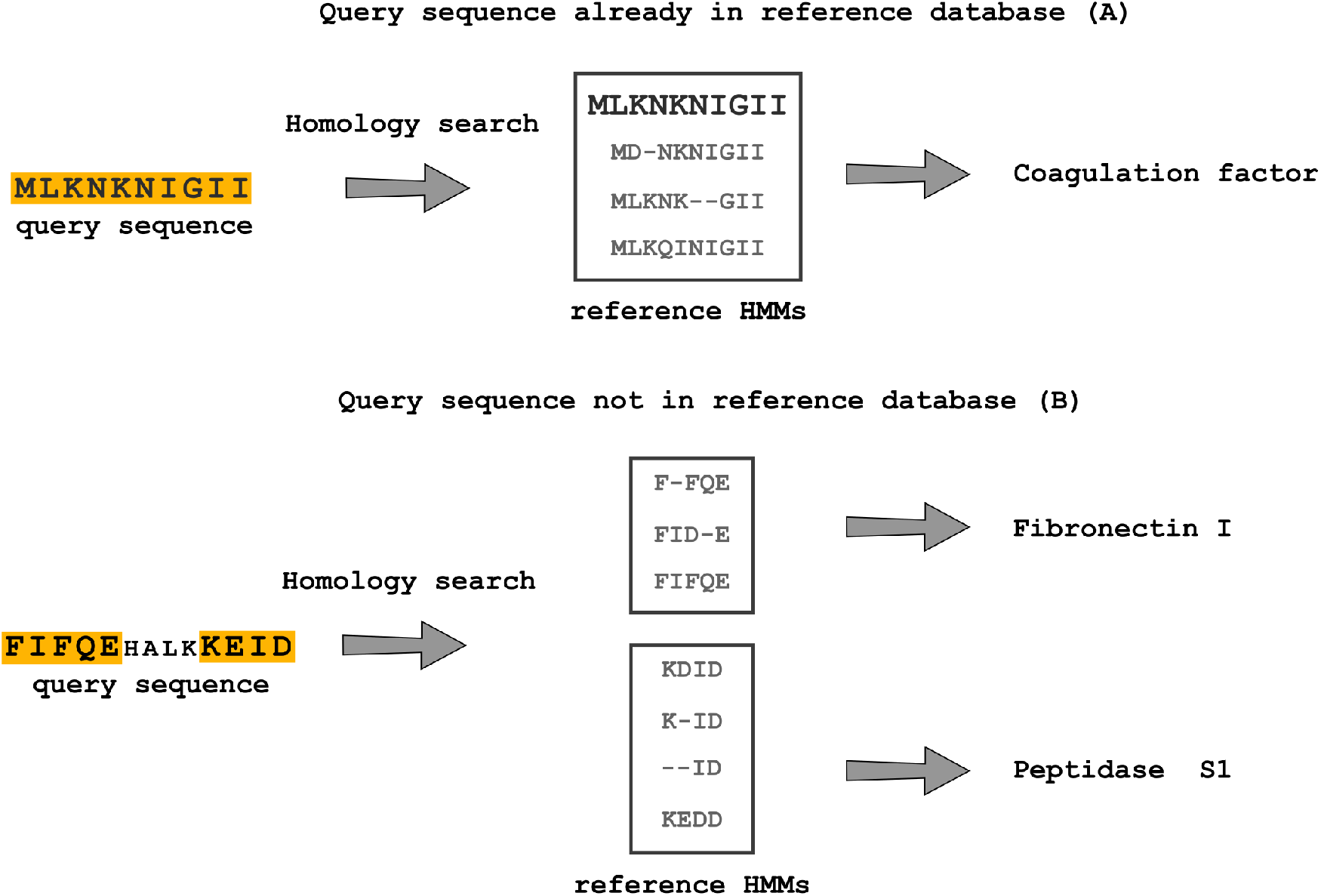
The impact of the reference datasets’ completeness on protein function annotation. While in the first scenario we can correctly predict the function of the protein sequence since it had been previously identified and included in the reference HMMs, the same is not possible in the second scenario as this protein sequence has not been previously described.

The use of eggNOG’s TSHMMs resulted in a 0.029 higher precision, however, this improvement, as seen in **Figure 3**, was not consistent across all the organisms. This may be due to a poorer quality of the TSHMMs for some organisms, which is both a consequence of the issues with the current taxonomy classification system [46, 45] and lack of knowledge regarding highly resolved taxa [12]. Model organisms such as *Escherichia coli* and *Saccharomyces cerevisiae* benefited the most from TSHMMs, both due to the fact that the reference datasets already contain data specific to these organisms and that functions of proteins within model organisms are better experimentally described. Conversely, non-model organisms are often only computationally annotated by association, contributing to a weaker reference annotation (which can be observed by the higher rate of potentially new annotations in these organisms in **Figure 3**). Nonetheless, while experimental evidence remains the gold standard, it is unfeasible to ignore the need for computational methods to infer function. While steps in this direction have been taken [29, 7], taxa-resolved PFA remains a challenge.

We benchmarked Mantis against another state-of-the-art PFA tool - eggNOG-mapper, and have shown that Mantis is both more precise (0.11 higher precision) and faster (6.6 times faster). Although Mantis’ default execution heavily relies on the eggNOG reference datasets, the use of additional reference datasets led to marked precision improvements over eggNOG-mapper. When Mantis ran without the eggNOG dataset, except for *Escherichia coli* and *Saccharomyces cerevisiae*, it achieved higher precision than eggNOG-mapper, proving that despite using much less data, Mantis did not suffer from substantial losses in precision. This attests to the quality of the various reference datasets used, showcasing as well the possibility of running Mantis on a personal computer (something that would be impossible with eggNOG’s prohibitive size). When running both tools with the same data, eggNOG-mapper had a higher precision (0.045), we hypothesize that even though both tools use the same data, the homology search methods used (Mantis uses HMMER and eggNOG-mapper uses Diamond) may lead to slightly different outputs. Nevertheless, Mantis’ flexibility in introducing multiple reference datasets and speed gives it unique advantages over similar tools. As we discussed, there is still room for improvement in the hit processing algorithm DFS, our text mining approach, and the addition of genomic context-based annotation methods. However, despite the existence of these challenges, we have clearly shown the unique advantages of using Mantis.

## Conclusion

By making use of the synergistic nature of differently sourced high-quality reference datasets, Mantis produces reliable homology-based annotations. By allowing for total customization of these reference datasets, Mantis is also extremely flexible, being easily integrated and adapted towards various research goals. In conclusion we have herein shown that Mantis addresses a number of the current PFA challenges, resulting in a PFA tool that is a fast, precise, flexible, and competitive tool.

## Methods

### Accessibility and Scaling

Mantis automatically sets up its reference data by downloading several reference datasets, and, when necessary, reformatting the data to a standardized format and downloading any relevant metadata. These reference datasets can be customized via a config file [54]. It also dynamically configures its execution depending on the resources available. A conda environment is provided for an easier setup, as well as extensive documentation [59].

Mantis splits most of the workflow into sub-tasks and subsequently parallelizes them by continuously releasing tasks to workers from a global queue (via Python’s multiprocessing module). During each main task of the annotation workflow workers are recruited (the number of workers depends on the available hardware), which will then execute and consume all the tasks within the queue. When a worker has finished its job, it will execute another task from the queue, until there are no more tasks to execute. If the queue is well balanced, minimal idle time (time spent waiting for workers to get a new task) can be achieved.

Parallelization is achieved both by splitting the sample and reference data into chunks. During setup, large reference datasets (more than 5000 HMM profiles) are split into smaller chunks, this enables parallelization and ensures each annotation sub-task takes approximately the same time. Large samples are pre-processed by splitting these into chunks (sample chunk size is dynamically calculated). If the sample has 200,000 or fewer sequences, sequences are distributed by their length among the different chunks, so that each chunk has approximately the same number of residues. If the sample has more than 200,000 sequences, then sequences are distributed to each chunk independently of their length (this alternative method is an efficiency safeguard). This two-fold splitting achieves quasi-optimal queue balancing. Posterior workflow steps are also parallelized where possible. Mantis is also scalable to metagenomes. For more details on performance see **Annotating metagenomes** in supplements.

### Reference data and customization

Mantis, by default, uses multiple high-quality HMM sources – eggNOG v5 [29], Pfam [21], Kofam [3], and TIGRfam [22] (these default HMMs can be partially or entirely removed). To find more meaningful homologs through taxa-specific annotation, Mantis uses TSHMMs, which were originally compiled by eggNOG. TSHMMs metadata was extracted from the eggNOG SQL database such that it is readily available during execution. Custom HMM sources can also be added by the user, metadata integration of these is also possible (an example is available in Mantis’ repository [55]). Since some sources are more specific than others, the user may also customize the weight given to each source during consensus generation [56].

HMM profiles often only possess an identifier respective to the database they were downloaded from, which may not directly provide any discernible information. Mantis, when necessary, ensures that the predictions from these HMMs are linked to their respective metadata. For future reference, while an HMM is an individual profile, Mantis compiles all related HMM profiles into a single file making it indexable by HMMER. Thus when a certain HMM source is mentioned it refers to the collection of related HMM profiles.

### Taxa-specific annotation

Taxa-specific annotation (TSA) uses the TSHMMs and unspecific HMM made available by eggNOG, TSA however works differently with respect to the annotation method of the other reference datasets. When given taxonomy information (either a taxa name or NCBI ID) the taxonomic lineage of the organism is computed (e.g. for *Escherichia coli* the lineage would be 2 - 1224 - 1236 - 91347 - 543 - 561 - 562). TSA starts by searching for homologs in the most resolved HMM (in this case for taxa 562, if it exists). All valid homologs (respecting the e-value threshold) are extracted for each query sequence, and unannotated sequences are compiled into an intermediate fasta file. A new round of homolog search then starts for the sequences in the intermediate fasta but now on the HMM one level above the previous HMM (in this case the HMM 561) such that valid homologs are again extracted. This cycle repeats until all query sequences have valid homologs or until there are no more HMMs to search for. If there are still sequences to annotate, then these homologs are searched for in the unspecific eggNOG dataset. If no taxonomy information is given then the homology search starts with the unspecific eggNOG HMM.

### Input and output

MANTIS accepts protein fasta files as input. For example, if running MANTIS for an *Escherichia coli* sample, the user would execute *python mantis run mantis -t sample.faa -od Escherichia coli*. If annotating a taxonomically unclassified sample the user would omit *-od*. Mantis can also annotate several samples at once, running *python mantis run mantis -t samples.tsv*, where the TSV file is a tab-separated file that contains the sample’s identifier, their path, and taxonomy information (an example file is available in Mantis’ repository [57]). Compressed fasta files and directory paths (with fasta files) are also accepted as input.

Mantis outputs, for each sample, three tab-separated files [58]: i) a raw output *output_annotation.tsv* with all the predictions, their e-value, and coordinates; ii) *integrated_annotation.tsv*, with all the predictions, coordinates, e-values, and their respective metadata and identifiers (e.g. KOs, EC numbers, free-text description, etc); and iii) the main output file *consensus_annotation.tsv*, with each query protein identifier and their respective consensus annotation from the different reference sources (e.g. Pfam). These files provide contextualized output in a format that is both human and machine-readable.

### Multiple predictions per protein

HMMER outputs a *domtblout* file [62], where each line corresponds to a hit/match between the reference dataset and the unknown protein sequence. The e-value within the HMMER command limits the available solution space to be analyzed in the posterior processing steps. Each hit, among other information, contains the coordinates where the query sequences matched with the reference HMM profiles and the respective confidence score (e-value). The annotation of a protein sequence with multiple hits is a nontrivial problem, requiring thus the implementation of a method for the **intra-HMM** processing of hits.

We designed a method that generates and evaluates all possible combinations of hits by applying the DFS algorithm [33]. This algorithm allows the traversal of a tree-structured search space (i.e. each node is a protein prediction), whilst pruning solutions that do not respect predefined constraints (i.e. overlapping protein predictions), backtracking from leaf to root until the possible solution space is exhausted. Our method generates all the possible combination hits with the following method: i. Get one hit from the collection of hits and define it as the combination root hit; ii. Check which other hits overlap up to 10% (default value) [86] with previous hits and select one to add to our current combination of hits; iii. Repeat step ii until no more hits can be added; iv. Repeat steps i-iii so that we loop over all the other hits and all possible combinations are generated. We used Cython [8] to speed up the DFS implementation. Cython is an optimising static compiler for the Python programming language, allowing the compiler to generate very efficient C code from Cython code, in this case, functioning as a wrapper for the DFS algorithm. The total amount of possible combinations is 2^*N*^ − *X* − 1 where *N* is the number of hits the protein sequence has, *X* the number of impossible combinations (combinations with overlapping hits), and *1* the empty combination. This number scales exponentially, as a result, this method is not always computationally feasible (e.g. the query sequence is very large and has many small-sized hits). In such a scenario, should the DFS algorithm running time exceed 60 seconds, Mantis employs the previously described “heuristic” algorithm [86], which scales linearly and outputs an unique combination of hits. After generating all the possible combinations, each combination is evaluated according to several scores:

- *combination coverage* – combinations of hits that cover a large percentage of the protein sequence are more significant. This metric corresponds to 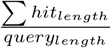.
- *combination e-value* – the combination of hits that have a lower sum of e-values are more significant. The e-value of each hit is scaled twice, once to reduce the range between different e-values (log10) and the second to understand how each hit e-value compares to the best/lowest hit e-value found for a certain sequence. The scaled e-values are then summed. This metric corresponds to Σ *MinMaxScale*(*log*10(*hit_e−value_*)).
- *amount of hits per combination* - combinations with few but large-sized hits are more significant, and vice-versa. This metric corresponds to *combo_length_*.

This *combined score* is defined by the following equation:

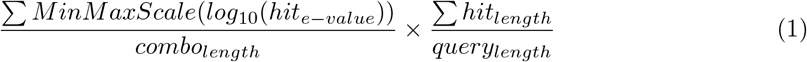

The combination with the highest *combined score* is then selected. Our implementation thus applies a two-fold quality control, initially by limiting the *domtblout* solution space and secondly by hierarchically ordering and selecting the most significant combination of hits.

### Using multiple reference datasets

An unannotated protein sequence may match with zero, one, or multiple reference HMM profiles, from one or more reference datasets. When a protein sequence has multiple predictions from different data sources it is necessary to identify complementary annotations, so that no information is lost. By linking the metadata respective to the HMM profiles to the now annotated protein sequence, we can identify similar annotations and integrate multiple reference datasets into one final consensus annotation. For the integration of multiple reference datasets, a two-fold text mining approach was used: 1. *Consensus between identifiers*; and 2. *Consensus between the free-text annotation description*.

Each reference data source includes metadata relevant to the HMM profiles herein; metadata may include multiple intra and/or inter database identifiers as well as free-text descriptions. Identifiers are extracted either through source-specific metadata parsing or by using regular expressions. Free-text descriptions are extracted by source-specific metadata parsing. When using custom data, metadata must have a standardized format [55].

The *consensus between identifiers* is calculated by identifying intersections between the different sources. Identifiers within the free-text annotation descriptions are extracted and used here. If no consensus between identifiers is found, then we proceed with a consensus calculation between annotation descriptions. The *consensus between free-text annotation descriptions* is done by comparing the free-text descriptions from two sources and evaluating whether these are similar. This step starts with parsing and pre-processing of the text within the annotation descriptions. It is followed by the lexical classification of the different processed words, so that irrelevant words are not considered in the next steps. This classification is done via a tagger [50] independent of context. This tagger uses the Wordnet lexicon [84] to identify the most common lexical category of any given word. Wordnet is used as it provides one of the most comprehensive lexical databases in the English language. To adjust this lexicon to biological data, gene ontologies [6, 74] are parsed, processed, and added as nouns to the tagger. If there are still untagged words, these are contextually classified with a pre-trained Perceptron tagger [61, 1]. Finally, the words tagged as determiners, pronouns, particles, or conjunctions are removed. Each word is then given a weight so that less specific words (e.g. “protein”) have a lower weight, and vice versa. We chose the metric Term Frequency-Inverse Document Frequency (*TF-IDF*) as it is a good metric for scoring how important a word is in a document (in the current context, documents are the annotations’ descriptions) relative to the entire corpus (reference collection of annotation descriptions), and has been successfully used in the past [9, 25, 73, 28]. We pre-calculated a frequency table of all the words from a collection of 561.911 reviewed proteins from Swiss-Prot [77] (as of 2020/04/14). With this frequency table, the IDF metric (corpus-wide/global weight of each word) can be calculated, such that words that appear in too many annotation descriptions in the corpus are less important. TF is a local metric specific to the annotation description being analysed, weighing words such that more frequent words in an annotation description are given a higher weight. Finally, *TF-IDF* is calculated with the following equation:

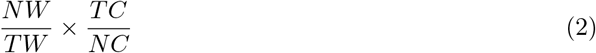

where *NW* is the amount of times a word appears in an annotation description, *TW* the total amount of words in an annotation description, *TC* the total amount of annotation descriptions in the corpus (561.911), and *NC* the total amount of times a certain word appears in the corpus. The *TF-IDF* score is then locally scaled so that we can better understand which words are more relevant within the current annotation description. After these steps we obtain a TF-IDF scaled vector for each annotation description. Finally, for similarity analysis, we calculate the cosine distance [23] between all the annotation description TF-IDF scaled vectors (which would come from different data sources), effectively measuring how similar these two vectors are. Should the words they contain and their importance within the annotation description be similar, the annotations are given a high similarity score. For a detailed description of how the similarity threshold was set see “**Defining a similarity threshold**” in the supplements. The calculation of the similarity of annotation descriptions is available as a standalone package [53]. In the context of the Mantis’ workflow, **inter-HMM** combinations of hits are generated and then expanded using the text mining method just described. Since several groups of consensus annotations may be generated, we evaluate their quality and select the best one, considering the following: percentage of the sequence covered by the hits in the consensus, the significance of the hits (e-value) in the consensus, significance of the reference datasets (customizable), and the number of different reference datasets in the consensus.

### Establishing a test environment

For annotation quality benchmarking we evaluate each annotation produced by Mantis and check whether it matches (True-positives) or not (False-positives) with the respective reference annotation. **True-Positives** (TP) are evaluated via two main methods: i) identifiers match and description match; the latter is detected via the same text mining approach used to generate the consensus annotations. Annotations which do not match with the reference annotation are classified as **False-Positives** (FP). Low quality annotations were removed (descriptions containing “unknown function”, “uncharacterized protein”, “hypothetical protein” or having Pfam’s “domain-unknown-function”/DUF identifiers) both from Mantis’ annotation as well as the reference annotations. Whenever Mantis outputs a valid annotation but the reference is either of poor quality or non-existent, then this annotation is considered a potentially **new annotation**. **Annotation coverage** is defined here as the number of Mantis’ annotations divided by the total amount of protein sequences in a sample. The **% TPs** and **% FPs** are, respectively, defined as the number of TPs and FPs divided by the number of sequences in the sample. **Precision** as 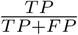 (ranging from 0 to 1), precision was chosen to represent annotation quality.

Finally, we benchmarked Mantis against another PFA tool - eggNOG-mapper. This tool was chosen as it shares many key properties with Mantis: it directly takes as input a protein sequence fasta, it uses a large and comprehensive reference database for homology search, and is capable of outputting taxa-resolved annotations. Unlike Mantis, eggNOG-mapper exclusively uses the eggNOG database, thus not allowing for customization or addition of new reference datasets; additionally, for homology search, Mantis uses HMMER [62] and eggNOG-mapper uses Diamond [11].

All tests ran on an HPC with Dell C6320, 2 * Intel Xeon E5-2680 v4 @ 2.4 GHz [78], each core had 4GB of RAM. Unless specified, all tests ran with 25 cores and 100GB RAM (actual minimum hardware requirements are much lower), in addition, the same methodology and nomenclature apply to any other benchmarked tools described in this paper. Mantis used HMMER v3.2.1. The local version of eggNOG-mapper is v1.0.32.0.1-14-gbf04860 with database v2.0.

Mantis uses HMMER, which runs via the following command:

`hmmsearch --domtblout output.domtblout -E e_value --notextw dataset.hmm sample.faa`

Mantis ran with the following command:

*python mantis run_mantis -t sample.faa -od “NCBI ID”*

eggNOG-mapper was executed with the following command:

*python emapper.py -i sample.faa -o output_folder -m diamond*

### Sample selection

As an initial testing dataset we started by downloading all the reviewed Swiss-Prot [77] protein entries created after 2010 (until 2020/04/14), along with their respective sequences, annotations, and annotations scores. We then split these entries by date, 2010-2020, 2015-2020,2018-2020, and 2020 only. For genomic sample benchmarking we selected organisms widely used in microbial community standards. The respective genomes, proteomes, and reference annotations were then downloaded from Uniprot on 2020/05/26 (supplemental **Table 3**). These genomic samples were also used for comparing Mantis to eggNOG-mapper.

### Testing different e-value thresholds

Different e-value thresholds were tested: 1*e*^−3^, 1*e*^−6^,1*e*^−9^,1*e*^−12^,1*e*^−15^,1*e*^−18^,1*e*^−21^,1*e*^−24^,1*e*^−27^,1*e*^−30^, and a dynamic threshold. The dynamic threshold was set according to the query sequence length, which has been previously shown to provide better results with BLAST [76]. For the dynamic threshold, for sequences with less than 150 amino acids, the e-value threshold was set to 1*e^−^*10, if above 150 and below 250, 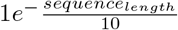, and if above 250, 1*e^−^*25. The Swiss-Prot dataset was then annotated with all the different e-value thresholds and each output was compared to the reference annotations.

### Availability of source code and requirements

- Project name: Mantis
- Project home page: https://github.com/PedroMTQ/mantis
- Operating system(s): Linux
- Programming language: Python
- Other requirements: Python 3+, HMMER 3+, and several Python packages (requests, numpy, nltk, sqlite, psutil, cython)
- License: MIT license at https://github.com/PedroMTQ/mantis/blob/master/LICENSE

### Availability of supporting data and materials

The code relative to the NLP analysis is available at https://github.com/PedroMTQ/nlp_annotations. The data and code supporting the results of this article are available https://github.com/PedroMTQ/mantis/wiki/Resources/mantis_data.7z. The supplemental pdf “supplements.pdf” contains information on how the similarity threshold was defined, the performance of metagenome annotation, efficiency benchmark, benchmark against other PFA tools and all the tables that were referenced in this paper. The non-averaged results supporting this paper are also available in the supplemental file “supplements.xlsx”.

## Supporting information

Supplemental tables raw

## Declarations

### List of abbreviations

DFS: Depth First Search
FP: False Positives
FN: alse Negatives
HMM: Hidden Markov Models
PFA: Protein Function Annotation
TF-IDF: Term Frequency-Inverse Document Frequency
TN: True Negatives
TP: True Positives
TSA: Taxa Specific Annotation
TSHMM: Taxa-Specific HMM

## Author’s Contributions

Pedro Queirós and Patrick May initiated the study. Mantis was developed and benchmarked by Pedro Queirós. Patrick May supervised the protein function annotation work. Francesco Delogu, Oskar Hickl, and Paul Wilmes gave feedback on the paper. All authors proof-read and approved of the content in this research paper.

## Acknowledgements

The experiments presented in this paper were carried out using the HPC facilities of the University of Luxembourg [78]. We would like to thank Tomila Litvishko for proof-reading this research paper. We would like to acknowledge all the creators of the reference datasets and tools used by Mantis, building upon the complementary knowledge of others truly takes the field forward.

## References

[1] A Good Part-of-Speech Tagger in about 200 Lines of Python. 2013.

[2] Stephen F. Altschul et al. “Basic local alignment search tool”. In: Journal of Molecular Biology 215.3 (1990), pp. 403–410. issn: 0022-2836. doi:10.1016/S0022-2836(05)80360-2.

[3] Takuya Aramaki et al. “KofamKOALA: KEGG Ortholog assignment based on profile HMM and adaptive score threshold”. In: Bioinformatics 36.7 (2020), pp. 2251–2252. issn: 1367-4803. doi:10.1093/bioinformatics/btz859. (Visited on 06/25/2020).

[4] Fabricio Almeida Araujo et al. “GO FEAT: a rapid web-based functional annotation tool for genomic and transcriptomic data”. In: Scientific Reports 8.1 (2018), p. 1794. issn: 2045-2322. doi:10.1038/s41598-018-20211-9.

[5] Carolina Arias et al. “KSHV 2.0: A Comprehensive Annotation of the Kaposi’s Sarcoma-Associated Herpesvirus Genome Using Next-Generation Sequencing Reveals Novel Genomic and Functional Features”. In: PLOS Pathogens 10.1 (2014), e1003847. issn: 1553-7374. doi:10.1371/journal.ppat.1003847.

[6] Michael Ashburner et al. “Gene Ontology: tool for the unification of biology”. In: Nature genetics 25.1 (2000), pp. 25–29. issn: 1061-4036. doi:10.1038/75556.

[7] Ramy K. Aziz et al. “The RAST Server: Rapid Annotations using Subsystems Technology”. In: BMC Genomics 9.1 (2008), p. 75. issn: 1471-2164. doi:10.1186/1471-2164-9-75.

[8] Stefan Behnel et al. “Cython: The Best of Both Worlds”. In: Computing in Science Engineering 13.2 (2011), pp. 31–39. issn: 1558-366X. doi:10.1109/MCSE.2010.118.

[9] Sidahmed Benabderrahmane et al. “IntelliGO: a new vector-based semantic similarity measure including annotation origin”. In: BMC Bioinformatics 11 (2010), p. 588. issn: 1471-2105. doi:10.1186/1471-2105-11-588.

[10] Karsten M. Borgwardt et al. “Protein function prediction via graph kernels”. In: Bioinformatics 21 (suppl 1 2005), pp. i47–i56. issn: 1367-4803. doi:10.1093/bioinformatics/bti1007.

[11] Benjamin Buchfink, Chao Xie, and Daniel H. Huson. “Fast and sensitive protein alignment using DIAMOND”. In: Nature Methods 12.1 (2015), pp. 59–60. issn: 1548-7105. doi:10.1038/nmeth.3176.

[12] Robin Buell et al. Breaking the Bottleneck of Genomes: Understanding Gene Function Across Taxa. US Department of Energy, Office of Biological and Environmental Research, 2018, p. 72.

[13] Agnes Chapel et al. “An Extended Proteome Map of the Lysosomal Membrane Reveals Novel Potential Transporters”. In: Molecular & Cellular Proteomics 12.6 (2013), pp. 1572–1588. issn: 1535-9476, 1535-9484. doi:10.1074/mcp.M112.021980.

[14] Nikolai Daraselia et al. “Automatic extraction of gene ontology annotation and its correlation with clusters in protein networks”. In: BMC bioinformatics 8 (2007), p. 243. issn: 1471-2105. doi:10.1186/1471-2105-8-243.

[15] Lei Deng et al. “MADOKA: an ultra-fast approach for large-scale protein structure similarity searching”. In: BMC Bioinformatics 20.19 (2019), p. 662. issn: 1471-2105. doi:10.1186/s12859-019-3235-1.

[16] Rezvan Ehsani and Finn Drabløs. “TopoICSim: a new semantic similarity measure based on gene ontology”. In: BMC bioinformatics 17.1 (2016), p. 296. issn: 1471-2105. doi:10.1186/s12859-016-1160-0.

[17] Diana Ekman et al. “Multi-domain Proteins in the Three Kingdoms of Life: Orphan Domains and Other Unassigned Regions”. In: Journal of Molecular Biology 348.1 (2005), pp. 231–243. issn: 0022-2836. doi:10.1016/j.jmb.2005.02.007.

[18] Kenneth W. Ellens et al. “Confronting the catalytic dark matter encoded by sequenced genomes”. In: Nucleic Acids Research 45.20 (2017), pp. 11495–11514. issn: 1362-4962. doi:10.1093/nar/gkx937.

[19] Iddo Friedberg. “Automated protein function prediction—the genomic challenge”. In: Briefings in Bioinformatics 7.3 (2006), pp. 225–242. issn: 1467-5463. doi:10.1093/bib/bbl004.

[20] Sonali Vijay Gaikwad, Archana Chaugule, and Pramod Patil. “Text mining methods and techniques”. In: International Journal of Computer Applications 85.17 (2014).

[21] Sara El-Gebali et al. “The Pfam protein families database in 2019”. In: Nucleic Acids Research 47 (D1 2019), pp. D427–D432. issn: 0305-1048. doi:10.1093/nar/gky995.

[22] Daniel H. Haft et al. “TIGRFAMs and Genome Properties in 2013”. In: Nucleic Acids Research 41 (Database issue 2013), pp. D387–D395. issn: 0305-1048. doi:10.1093/nar/gks1234.

[23] Jiawei Han, Jian Pei, and Micheline Kamber. Data mining: concepts and techniques. Elsevier, 2011. isbn: 0-12-381480-4.

[24] Geoffrey D Hannigan et al. “A deep learning genome-mining strategy for biosynthetic gene cluster prediction”. In: Nucleic Acids Research 47.18 (Aug. 2019), e110–e110. issn: 0305-1048. doi:10.1093/nar/gkz654.

[25] Miao Hao and Ke Fan. “A Method for Calculating the Similarity of TF - IDF Texts for Synonyms in Biomedical Domains”. In: 2017 5th International Conference on Frontiers of Manufacturing Science and Measuring Technology (FMSMT 2017). Atlantis Press, 2017, pp. 578–583. isbn: 978-94-6252-331-9. doi:10.2991/fmsmt-17.2017.117.

[26] Anna Heintz-Buschart et al. “Integrated multi-omics of the human gut microbiome in a case study of familial type 1 diabetes”. In: Nature Microbiology 2.1 (2016). Number: 1 Publisher: Nature Publishing Group, pp. 1–13. issn: 2058-5276. doi:10.1038/nmicrobiol.2016.180.

[27] Chung-Chi Huang and Zhiyong Lu. “Community challenges in biomedical text mining over 10 years: success, failure and the future”. In: Briefings in Bioinformatics 17.1 (2016), pp. 132–144. issn: 1477-4054. doi:10.1093/bib/bbv024.

[28] Yue Huang, Mingxin Gan, and Rui Jiang. “Ontology-Based Genes Similarity Calculation with TF-IDF”. In: vol. 7473. 2012, pp. 600–607. doi:10.1007/978-3-642-34062-8_78.

[29] Jaime Huerta-Cepas et al. “eggNOG 5.0: a hierarchical, functionally and phylogenetically annotated orthology resource based on 5090 organisms and 2502 viruses”. In: Nucleic Acids Research 47 (D1 2019), pp. D309–D314. issn: 0305-1048. doi:10.1093/nar/gky1085.

[30] Jaime Huerta-Cepas et al. “Fast Genome-Wide Functional Annotation through Orthology Assignment by eggNOG-Mapper”. In: Molecular Biology and Evolution 34.8 (2017), pp. 2115–2122. issn: 0737-4038. doi:10.1093/molbev/msx148.

[31] Massimo Iorizzo et al. “De novo assembly and characterization of the carrot transcriptome reveals novel genes, new markers, and genetic diversity”. In: BMC Genomics 12.1 (2011), p. 389. issn: 1471-2164. doi:10.1186/1471-2164-12-389.

[32] Philip Jones et al. “InterProScan 5: genome-scale protein function classification”. In: Bioinformatics 30.9 (2014), pp. 1236–1240. issn: 1367-4803. doi:10.1093/bioinformatics/btu031.

[33] Navneet Kaur and Deepak Garg. “Analysis of the Depth First Search Algorithms”. In: Data mining and knowledge engineering 4 (2012), pp. 37–41.

[34] Kevin P. Keegan, Elizabeth M. Glass, and Folker Meyer. “MG-RAST, a Metagenomics Service for Analysis of Microbial Community Structure and Function”. In: Methods in Molecular Biology (Clifton, N.J.) 1399 (2016), pp. 207–233. issn: 1940-6029. doi:10.1007/978-1-4939-3369-3_13.

[35] William Klimke et al. “Solving the Problem: Genome Annotation Standards before the Data Deluge”. In: Standards in Genomic Sciences 5.1 (2011), pp. 168–193. issn: 1944-3277. doi:10.4056/sigs.2084864.

[36] Michael Kramer et al. “Inferring gene ontologies from pairwise similarity data”. In: Bioinformatics (Oxford, England) 30.12 (2014), pp. i34–42. issn: 1367-4811. doi:10.1093/bioinformatics/btu282.

[37] Jonathan G. Lees et al. “Gene3D: Multi-domain annotations for protein sequence and comparative genome analysis”. In: Nucleic Acids Research 42 (Database issue 2014), pp. D240–D245. issn: 0305-1048. doi:10.1093/nar/gkt1205.

[38] Meng Liu and Paul D. Thomas. “GO functional similarity clustering depends on similarity measure, clustering method, and annotation completeness”. In: BMC bioinformatics 20.1 (2019), p. 155. issn: 1471-2105. doi:10.1186/s12859-019-2752-2.

[39] Marc Lohse et al. “Mercator: a fast and simple web server for genome scale functional annotation of plant sequence data”. In: Plant, Cell & Environment 37.5 (2014), pp. 1250–1258. issn: 1365-3040. doi:10.1111/pce.12231.

[40] Olivia U. Mason et al. “Metagenomics reveals sediment microbial community response to Deepwater Horizon oil spill”. In: The ISME Journal 8.7 (2014), pp. 1464–1475. issn: 1751-7370. doi:10.1038/ismej.2013.254.

[41] Konstantinos Mavromatis et al. “Gene Context Analysis in the Integrated Microbial Genomes (IMG) Data Management System”. In: PLOS ONE 4.11 (2009), e7979. issn: 1932-6203. doi:10.1371/journal.pone.0007979.

[42] Alex L. Mitchell et al. “MGnify: the microbiome analysis resource in 2020”. In: Nucleic Acids Research 48 (D1 2020), pp. D570–D578. issn: 0305-1048. doi:10.1093/nar/gkz1035.

[43] Michael A. Mooney et al. “Functional and Genomic Context in Pathway Analysis of GWAS Data”. In: 30.9 (2014), pp. 390–400. issn: 0168-9525. doi:10.1016/j.tig.2014.07.004.

[44] Ross Overbeek et al. “The subsystems approach to genome annotation and its use in the project to annotate 1000 genomes”. In: Nucleic Acids Research 33.17 (2005), pp. 5691–5702. issn: 1362-4962. doi:10.1093/nar/gki866.

[45] Donovan H. Parks et al. “A complete domain-to-species taxonomy for Bacteria and Archaea”. In: Nature Biotechnology 38.9 (2020), pp. 1079–1086. issn: 1546-1696. doi:10.1038/s41587-020-0501-8.

[46] Donovan H. Parks et al. “A standardized bacterial taxonomy based on genome phylogeny substantially revises the tree of life”. In: Nature Biotechnology 36.10 (2018), pp. 996–1004. issn: 1546-1696. doi:10.1038/nbt.4229.

[47] Edoardo Pasolli et al. “Extensive Unexplored Human Microbiome Diversity Revealed by Over 150,000 Genomes from Metagenomes Spanning Age, Geography, and Lifestyle”. In: Cell 176.3 (2019), 649–662.e20. issn: 0092-8674. doi:10.1016/j.cell.2019.01.001.

[48] Jiajie Peng et al. “Measuring semantic similarities by combining gene ontology annotations and gene co-function networks”. In: BMC bioinformatics 16 (2015). issn: 1471-2105. doi:10.1186/s12859-015-0474-7.

[49] Catia Pesquita et al. “Semantic similarity in biomedical ontologies”. In: PLoS computational biology 5.7 (2009), e1000443. issn: 1553-7358. doi:10.1371/journal.pcbi.1000443.

[50] Slav Petrov, Dipanjan Das, and Ryan McDonald. “A Universal Part-of-Speech Tagset”. In: arXiv:1104.2086 [cs] (2011). (Visited on 06/25/2020).

[51] Friedhelm Pfeiffer and Dieter Oesterhelt. “A Manual Curation Strategy to Improve Genome Annotation: Application to a Set of Haloarchael Genomes”. In: Life 5.2 (2015), pp. 1427–1444. issn: 2075-1729. doi:10.3390/life5021427.

[52] Vasilis J. Promponas, Ioannis Iliopoulos, and Christos A. Ouzounis. “Annotation inconsistencies beyond sequence similarity-based function prediction – phylogeny and genome structure”. In: Standards in Genomic Sciences 10 (2015). issn: 1944-3277. doi:10.1186/s40793-015-0101-2.

[53] Pedro Queirós. Consensus between annotations. https://github.com/PedroMTQ/nlp_annotations. 2020.

[54] Pedro Queirós. Mantis - configuration file. https://github.com/PedroMTQ/mantis/blob/master/MANTIS.config. 2020.

[55] Pedro Queirós. Mantis - Custom HMMs. https://github.com/PedroMTQ/mantis/wiki/Configuration#custom-hmms. 2020.

[56] Pedro Queirós. Mantis - Custom HMMs weights. https://github.com/PedroMTQ/mantis/wiki/Configuration#setting-hmms-weight. 2020.

[57] Pedro Queirós. Mantis - Multiple samples. https://github.com/PedroMTQ/mantis/blob/master/tests/test_file.tsv. 2020.

[58] Pedro Queirós. Mantis - Output files. https://github.com/PedroMTQ/mantis/wiki/Output. 2020.

[59] Pedro Queirós. Mantis - Wiki. https://github.com/PedroMTQ/mantis/wiki. 2020.

[60] Pedro Queirós. Mantis: flexible and consensus-driven genome annotation. https://github.com/PedroMTQ/mantis. 2020.

[61] Frankie Roberston. Averaged perceptron tagger. 2016.

[62] Sean Roberts Eddy. HMMER. HMMER: biosequence analysis using profile hidden Markov models. 2020.

[63] Jae Yong Ryu, Hyun Uk Kim, and Sang Yup Lee. “Deep learning enables high-quality and high-throughput prediction of enzyme commission numbers”. In: Proceedings of the National Academy of Sciences 116.28 (2019), pp. 13996–14001. issn: 0027-8424, 1091-6490. doi:10.1073/pnas.1821905116.

[64] Alexandra M. Schnoes et al. “Annotation Error in Public Databases: Misannotation of Molecular Function in Enzyme Superfamilies”. In: PLoS Computational Biology 5.12 (2009). issn: 1553-734X. doi:10.1371/journal.pcbi.1000605.

[65] Torsten Seemann. “Prokka: rapid prokaryotic genome annotation”. In: Bioinformatics 30.14 (2014), pp. 2068–2069. issn: 1367-4803. doi:10.1093/bioinformatics/btu153. (Visited on 06/25/2020).

[66] Nicola Segata et al. “Computational meta’omics for microbial community studies”. In: Molecular Systems Biology 9 (May 14, 2013). issn: 1744-4292. doi:10.1038/msb.2013.22.

[67] Christian J. A. Sigrist et al. “New and continuing developments at PROSITE”. In: Nucleic Acids Research v41 (Database issue 2013), pp. D344–347. issn: 1362-4962. doi:10.1093/nar/gks1067.

[68] Luke T Slater et al. “Improved characterisation of clinical text through ontology-based vocabulary expansion”. In: bioRxiv (2020), p. 2020.07.10.197541. doi:10.1101/2020.07.10.197541.

[69] “Standardizing data”. In: Nature Cell Biology 10.10 (2008), pp. 1123–1124. issn: 1476-4679. doi:10.1038/ncb1008-1123.

[70] Martin Steinegger et al. “HH-suite3 for fast remote homology detection and deep protein annotation”. In: BMC Bioinformatics 20.1 (2019), p. 473. issn: 1471-2105. doi:10.1186/s12859-019-3019-7.

[71] Ahmet Sureyya Rifaioglu et al. “DEEPred: Automated Protein Function Prediction with Multi-task Feed-forward Deep Neural Networks”. In: Scientific Reports 9.1 (2019), p. 7344. issn: 2045-2322. doi:10.1038/s41598-019-43708-3.

[72] Damian Szklarczyk et al. “STRING v11: protein-protein association networks with increased coverage, supporting functional discovery in genome-wide experimental datasets”. In: Nucleic Acids Research 47 (D1 2019), pp. D607–D613. issn: 1362-4962. doi:10.1093/nar/gky1131.

[73] Georgi Tancev. Mining and Classifying Medical Documents. Medium. Library Catalog: towardsdatascience.com. 2019.

[74] “The Gene Ontology Resource: 20 years and still GOing strong”. In: Nucleic Acids Research 47 (D1 2019), pp. D330–D338. issn: 0305-1048. doi:10.1093/nar/gky1055.

[75] Paul Thompson et al. “The BioLexicon: a large-scale terminological resource for biomedical text mining”. In: BMC Bioinformatics 12.1 (2011), p. 397. issn: 1471-2105. doi:10.1186/1471-2105-12-397.

[76] Michelle L. Treiber et al. “Pre-and post-sequencing recommendations for functional annotation of human fecal metagenomes”. In: BMC Bioinformatics 21.1 (2020), p. 74. issn: 1471-2105. doi:10.1186/s12859-020-3416-y.

[77] “UniProt: a worldwide hub of protein knowledge”. In: Nucleic Acids Research 47 (D1 2019), pp. D506–D515. issn: 0305-1048. doi:10.1093/nar/gky1049.

[78] Sébastien Varrette et al. “Management of an Academic HPC Cluster: The UL Experience”. In: (2014). url: https://hpc.uni.lu.

[79] Alexei Vazquez et al. “Global protein function prediction from protein-protein interaction networks”. In: Nature Biotechnology 21.6 (2003), pp. 697–700. issn: 1546-1696. doi:10.1038/nbt825.

[80] Sheng Wang et al. “Annotating gene sets by mining large literature collections with protein networks”. In: Pacific Symposium on Biocomputing. Pacific Symposium on Biocomputing 23 (2018), pp. 602–613. issn: 2335-6936.

[81] Leon Weber et al. “HunFlair: An Easy-to-Use Tool for State-of-the-Art Biomedical Named Entity Recognition”. In: arXiv preprint arXiv:2008.07347 (2020).

[82] James C. Whisstock and Arthur M. Lesk. “Prediction of protein function from protein sequence and structure”. In: Quarterly Reviews of Biophysics 36.3 (2003), pp. 307–340. issn: 0033-5835. doi:10.1017/s0033583503003901.

[83] Mark D. Wilkinson et al. “The FAIR Guiding Principles for scientific data management and stewardship”. In: Scientific Data 3.1 (2016), p. 160018. issn: 2052-4463. doi:10.1038/sdata.2016.18.

[84] WordNet — A Lexical Database for English. 2010.

[85] Sitao Wu et al. “WebMGA: a customizable web server for fast metagenomic sequence analysis”. In: BMC genomics 12 (2011). issn: 1471-2164. doi:10.1186/1471-2164-12-444.

[86] Corin Yeats, Oliver C. Redfern, and Christine Orengo. “A fast and automated solution for accurately resolving protein domain architectures”. In: Bioinformatics 26.6 (2010), pp. 745–751. issn: 1367-4803. doi:10.1093/bioinformatics/btq034.

[87] Zhiqiang Zeng et al. “Survey of Natural Language Processing Techniques in Bioinformatics”. In: Computational and Mathematical Methods in Medicine 2015 (2015), p. 674296. issn: 1748-6718. doi:10.1155/2015/674296.

[88] Bihai Zhao et al. “An efficient method for protein function annotation based on multilayer protein networks”. In: Human Genomics 10 (2016). issn: 1473-9542. doi:10.1186/s40246-016-0087-x.

